# Renal Localization of *Plasmodium vivax* and Zoonotic Monkey Parasite *Plasmodium knowlesi* Derived Components in Malaria Associated Acute Kidney Injury

**DOI:** 10.1101/544726

**Authors:** Pragyan Acharya, Atreyi Pramanik, Charandeep Kaur, Kalpana Kumari, Rajendra Mandage, Amit Kumar Dinda, Jhuma Sankar, Arvind Bagga, Sanjay Kumar Agarwal, Aditi Sinha, Geetika Singh

## Abstract

**Objectives:** Acute kidney injury (AKI) is a frequent clinical manifestation of severe malaria. Although the pathogenesis of *P. falciparum* severity is attributed to cytoadherence, the pathogenesis due to *P. vivax* and *P. knowlesi* are not very well understood and their presence within the host tissue has not been demonstrated. Therefore, the objective of this study was to determine if *P. vivax* and *P. knowlesi* interact with host renal tissue in malarial AKI.

**Methods:** Archival FFPE renal biopsies from 16 patients having AKI with suspected or confirmed malaria diagnosis, and 5 patients with non-malarial AKI, were subjected to histopathological analysis and DNA extraction. The total DNA from these renal biopsies was used to investigate the presence of the 5 species of human malaria parasite: *P. falciparum, P. vivax, P. knowlesi, P. malariae* and, *P. ovale*. The results of histopathology and PCR analysis were analyzed to understand the pathogenesis of malaria associated AKI.

**Results:** Renal biopsies from malarial AKI patients were found to harbor DNA from *P. vivax* (14 of 16), the zoonotic monkey parasite *P. knowlesi* (7) and *P. falciparum* (5) in the form of mono-infections as well as mixed infections. Sanger sequencing was used to confirm species identity. Histopathological analysis revealed the presence of microvascular injury as a common feature of *P. vivax* and *P. knowlesi* related AKI. Hemozoin could be detected within the renal tissue, which indicated the presence of metabolically active parasites.

**Conclusions:** *P. vivax* and *P. knowlesi* associated AKI may involve interaction of the host renal tissue with parasite derived components such as *Plasmodium* DNA and hemozoin which may lead to innate immune activation and host tissue injury.

## INTRODUCTION

Human malaria is known to be caused by five species of *Plasmodium* parasites-*P. falciparum, P. vivax, P. knowlesi, P. ovale* and *P. malariae*. While *P. falciparum* is found all over the tropical world, *P. vivax* is highly prevalent in non-African regions such as S. America, India, S. E. Asia, and *P. knowlesi* has been shown to cause malaria predominantly in S.E. Asia [1–4] Infection with *Plasmodium* species can manifest as severe malaria in which fever is accompanied by extreme outcomes such as organ dysfunction, cerebral malaria, and coma; or as mild malaria, which presents as chills, and fever accompanied by milder symptoms. Acute kidney injury (AKI), is a frequently presentation in severe malaria due to both *P. falciparum* and *P. vivax*, and more recently has also been shown to be associated with *P. knowlesi* [5–8]. In fact, overall AKI has been shown to have a global prevalence in about 20-50% hospitalized malaria cases [9]. Pathogenesis of *P. falciparum* is attributed to its cytoadherence to endothelial cells of tissue capillaries. *P.knowlesi*-associated severe malaria, specifically AKI, is hypothesized to be due to the haemolysis of RBCs and subsequent release of cell-free haemoglobin leading to oxidative damage [10]. Additionally, *P. knowlesi ex vivo* cultures have been shown to bind to host endothelial receptors ICAM-1 and VCAM [11]. Mechanisms of *P. vivax* severity are unclear although *P. vivax* has been shown to adhere to host receptors with low affinity and recent studies implicate infected RBC phosphatidylserine in host cytoadherence [12]. However, unlike *P. falciparum*, which has been shown to associate with host tissues, neither *P. knowlesi* nor *P. vivax* have ever been localized to any host tissues including the kidney. Therefore the pathogenesis of AKI as well asother aspects of severe malaria due to *P. vivax* and *P. knowlesi* infections are not understood. It is important to study the molecular pathogenesis of *P. vivax* because of its high prevalence in the tropical world and its unexplained ability to cause severe disease. It is important to understand *P. knowlesi* pathogenesis since it is a rapidly emerging zoonotically transmitted parasite with an unknown biology with unknown molecular effectors. Therefore, in this study we have investigated if *P. vivax*es as well as the other four *Plasmodium* species are present within host renal tissue in malarial AKI. In order to investigate this we have conducted a retrospective study in renal biopsies of malaria associated-AKI patients at AIIMS, New Delhi, India. We report our observations here.

## METHODS

### Study design

This study was a cross-sectional retrospective analysis carried out as a collaboration between the Departments of Biochemistry, Pathology, Pediatrics and Nephrology at All India Institute of Medical Sciences (AIIMS), New Delhi, India. The ethics committee of the All India Institute of Medical Sciences, New Delhi has approved this study.

### Renal tissue samples

Archival formalin-fixed paraffin-embedded (FFPE) renal tissue blocks from 2011-2018 were retrieved from the Department of Pathology, AIIMS, New Delhi, India.

#### Cases

AKI patients presented with abdominal pain, fever and low urine output with suspected or confirmed malaria diagnosis.

#### Controls

Renal biopsies from AKI cases showing acute tubular injury or necrosis (ATN) and/or acute cortical necrosis (ACN) without clinical, laboratory, or histopathological evidence of malaria. Details of controls are presented in Supplementary table S1.

The individuals who performed the experiments were blinded to the identity of the controls and cases until the PCR outcomes had been obtained and documented.

Initial evaluation of patients included urine analysis, ultrasound, complete blood counts and measurements of creatinine, urea, electrolytes, pH and bicarbonate. Renal biopsies were processed for light microscopy by standard techniques. Diagnosis of malaria was based on peripheral blood smears.

### Histological examination

Histological sections of all FFPE tissues were stained with hematoxylin, periodic acid Schiff (PAS) and Jones methenamine silver stains. The glomeruli, tubules, interstitium, and blood vessels were examined in all the cores. To confirm that the crystals observed in the sections were hemozoin, a saturated solution of picric acid in ethanol was used (malarial bleach).

### Detection of Malaria parasites

a. Original diagnosis of malaria was based on detection of different parasitic forms of *Plasmodium* species in the peripheral blood smear (thick and thin smears stained by Giemsa). Species identification was done on thin film microscopy. In the cases where peripheral blood smear did not yield any parasites, diagnosis was carried out by physicians based on clinical presentation.
b. For detection of malaria parasite from renal tissue, DNA was extracted using the QiagenQiAmp DNA FFPE tissue kit as per the manufacturer’s protocol (Supplementary file). PCR amplification of extracted DNA for all the five *Plasmodium* species (*P. falciparum, P. vivax, P. knowlesi, P. malariae*and*P. ovale*) was carried out. The primers used for PCR amplification are listed in Supplementary Table S2. The PCR products were subjected to Sanger sequencing (Bencos Research Solutions Pvt. Ltd.) in order to confirm the species identity (Supplementary Table S3).

## RESULTS

In the present study, 16 FFPE samples with malaria associated AKI and5 control samples (AKI with non-malarial etiology) collected from 2011 to 2018 were included (Table1, Supplementary Table S1). Of the 16 cases, 12 presented with fever and associated oliguria, 1 presented with fever and associated hematuria; 1 as hematuria alone; 1 presented as AKI about one and half month after diagnosis as complicated *P. vivax* malaria; and 1 had presented with placental abruption associated with AKI (Table 1; column 3). The various histological diagnoses in these cases were acute cortical necrosis (ACN), acute tubular injury/necrosis (ATI/ATN), thrombotic microangiopathy (TMA), and interstitial fibrosis and tubular atrophy (IFTA). Of the 16 cases, 14 had evidence of cortical necrosis, TMA, or mesangiolysis, indicating vascular involvement (Table 1). The representative histopathology of cortical and tubular damage has been depicted in Supplementary Figure S1.

**Table 1.**
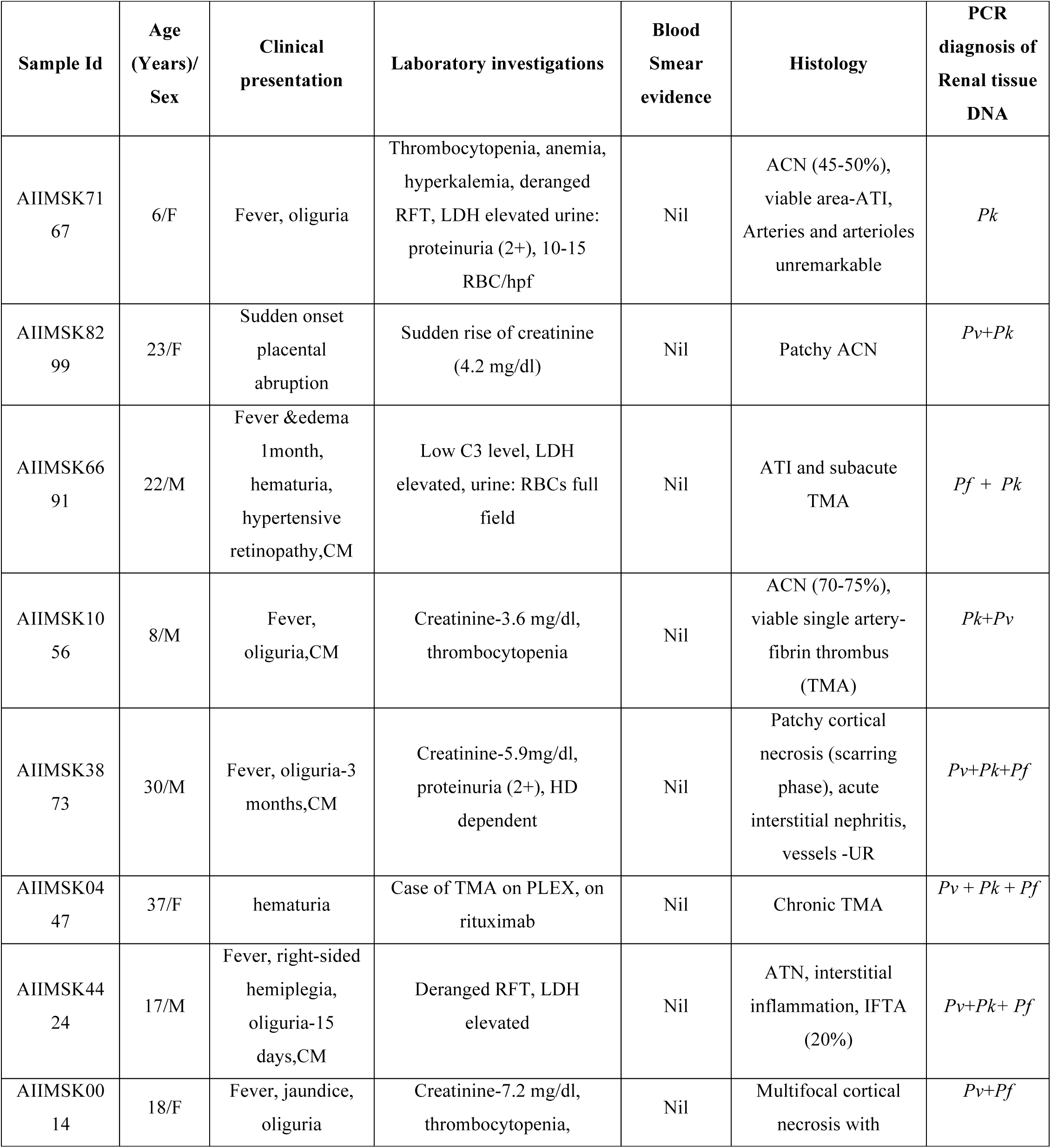
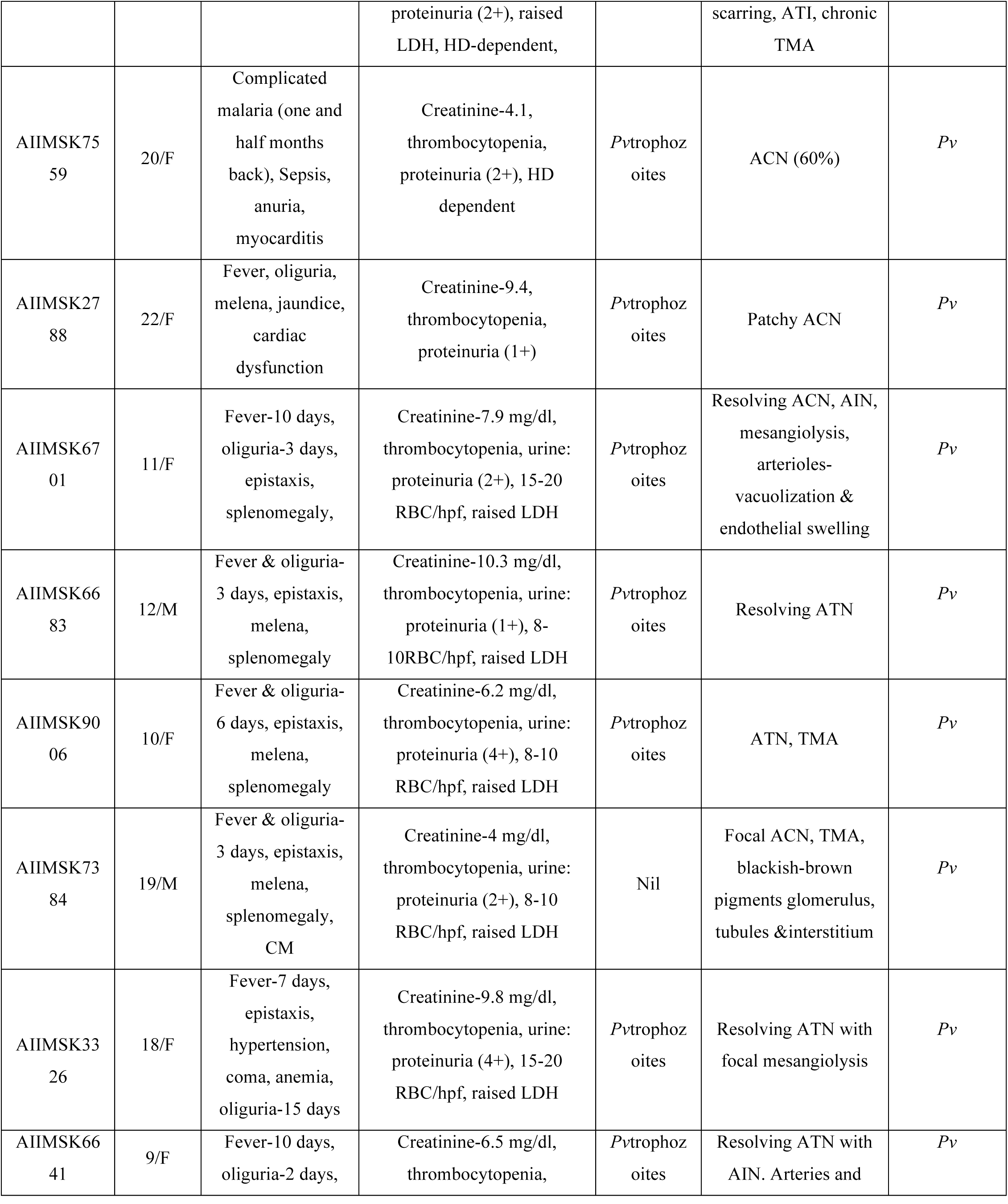
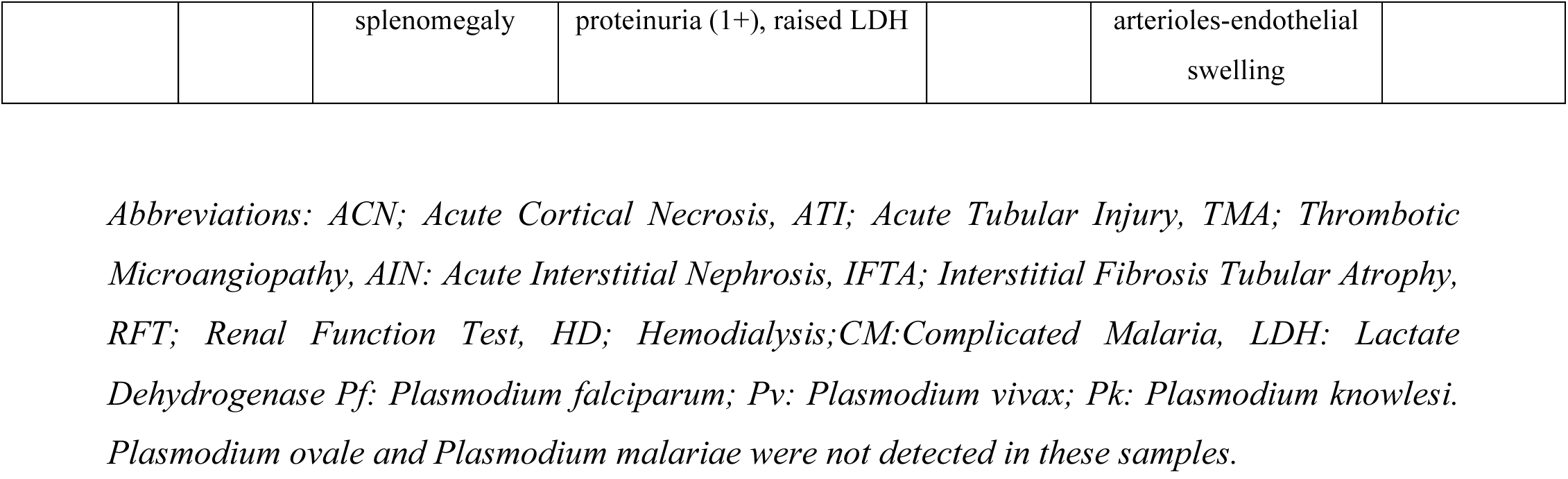
Summary of clinical presentation of AKI in patients with malaria infection with PCR diagnosis of the five *Plasmodium* species infecting human

The peripheral blood smears of 8caseswerefound to have infection of *P. vivax*alone, and 1 showed the presence of both *P. vivax* and *P. falciparum*trophozoites; 7 cases had no evidence ofmalaria from peripheral blood smears (Table 1, Column 5). Of these 7 samples, 5were clinically diagnosed as complicated malaria (CM), 1 presented with hematuria and 1 with placental abruption (Table 1). None of the control samples had any symptoms resembling malaria or any evidence of malaria from peripheral blood smear. They presented with ACN and TMA associated with non-malaria cases as described in Supplementary Table S1.

All the 16 cases and 5 controls were subjected to PCR analysis for all the 5 *Plasmodium* species (*P. falciparum, P. vivax, P. knowlesi, P. malariae* and *P. ovale*). Among the 16 cases, 8 cases were found to have *P. vivax*, 1 had P. *knowlesi*, 1 had mixed infection of *P. vivax* and *P. falciparum*, 2 had mixed infections of *P. vivax* and *P. knowlesi*, 1 had mixed infection of *P. falciparum* and *P. knowlesi*, and 3 had mixed infections consisting of three parasites *P. vivax, P. falciparum*, and *P. knowlesi* (Table 1). Overall, of 16 cases with malaria associated AKI, the renal tissue of 14 had *P. vivax*, 7 had *P. knowlesi* and, 5 had *P. falciparum* infection. Interestingly, of the 7 cases of malarial AKI having *P. knowlesi* by PCR, all lacked peripheral smear evidence of the parasite (Table 1) but 4 presented as CM, 1had placental abruption with AKI, and 1 presented with TMA and required plasmapheresis (Table 1, Column 3 and 4). The *P. knowlesi* PCR products were Sanger sequenced followed by NCBI BLAST in order to confirm species identity (Table S3).

None of the control FFPE renal biopsies contained parasitic DNA by PCR analysis (Supplementary Table S1).

The origin of all malaria associated AKI cases were mapped to Delhi and Uttar Pardesh (Supplementary Fig S2).

## DISCUSSION

Overall, of 16 malaria cases presenting with AKI, 14 (∼88%) had *P. vivax*, 7 had *P. knowlesi* and 5 had *P. falciparum* infection as mono-infections as well as mixed-infections. Although several studies show association of AKI with peripheral blood *P. vivax and P. knowlesi* infection [5,6,13–15] we demonstrate direct tissue presence of the parasite derived DNA for both these species. Interestingly, of the 16 cases, 9 had no evidence of the parasite from microscopic examination of peripheral blood smears, suggesting that microscopic examination alone may not be sufficient for detection and diagnosis of *P. vivax and P. knowlesi*.

In addition to PCR evidence of the presence of parasite DNA, we found specific staining for iron containing hemozoin, which was completely absent in all the control samples. Hemozoin pigment is known to be produced only by replicating parasites [16]. When in circulation, it is rapidly phagocytosed by macrophages and dendritic cells and therefore, has a short circulating half life[17, 18] Hemozoin pigment in renal tissue has been demonstrated in the case of *P. falciparum* associated AKI [9]. Therefore, presence of hemozoin staining specifically in renal tubules and interstitiumis suggestive of the presence of replicating parasites at the site, rather than non specific accumulation of hemozoin. However, our data do not include evidence of direct molecular interactions between *P. vivax* and *P. knowlesi*, and the host renal tissue.

It is well recognized now that *P. vivax* is capable of causing complicated malaria and is associated with several syndromes such as anemia, thrombocytopenia, impaired liver and renal function, and altered sensorium, among other features[19–22]. However its pathogenesis is not clear. In *P. falciparum*, pathogenesis of complicated malaria is linked to tissue cytoadhesion of the parasite. However, unlike *P. falciparum*, the cytoadhesion of *P. vivax* is still under investigation. It is not clear if it interacts with tissue capillaries or localizes within tissues. Several studies have explored the potential of *P. vivax* to cytoadhere to host endothelial cells and have found *in vitro* evidence of low affinity binding [12,23,24]. We have addressed the question of tissue localization of *P. vivax* and the other four human malaria parasites in malaria-associated acute kidney injury (AKI). The data presented here provide evidence for the renal presence of *P. vivax* derived DNA and hemozoin. Surprisingly we also found DNA originating from *P. knowlesi*, the zoonotic parasite known to infect Rhesus macaques (*Macaca mulatta*), within the renal tissue of many malaria AKI cases (7 of 16).

This is the first study to demonstrate presence of *P. knowlesi* derived DNA in the renal tissue. This is an important observation since the response of *P. knowlesi* to current anti-malarial therapy is not well understood [14,15] and may affect treatment outcomes in AKI cases that have co-infections of *P. vivax* and *P. knowlesi* within the renal tissue. This study also sheds light on potential pathogenesis of *P. vivax* and *P. knowlesi* related malaria severity. In the absence of evidence to show direct pathogen-host intercellular contact, at the very least, this study shows that parasite DNA, and hemozoin indicative of actively replicating parasites, are present in host renal tissue. Parasite DNA and hemozoin have both been shown to act as pathogen associated molecular patterns (PAMPS) that initiate TLR 7, TLR 9 or cytosolic DNA receptor STINGmediated activation of innate immune cells [26,27]. It is possible that irrespective of the parasite cytoadherence, *P. vivax* and *P. knowlesi* might be causing renal tissue damage through innate immune inflammation activated by PAMPs such as parasite dsDNA, hemozoin and membrane casts.

It is interesting to note that many of the cases harboring renal *P. knowlesi* were from New Delhi, and Uttar Pradesh where there has been a reported explosion of *Rhesus macaque* population in recent times (Supplementary Fig S2) [28]. This highlights the importance of malaria elimination not only in humans and mosquito vectors, but also in other hosts living in human proximity.

The major limitation of this study is the lack of microscopic evidence to investigate the presence of parasite cells within host renal tissue. Since a majority of these samples were archival, and taken predominantly for histopathological evaluation, their remains were limited in quantity and could only be repurposed for any one application-DNA isolation or microscopy. We have utilized them for DNA isolation to ensure nucleic acid based confirmation for many of the suspected malaria cases where microscopic evidence from peripheral smears were not available and they were categorized as malaria cases based on the clinical presentations. The major strength of our study is that it provides the first evidence of parasite derived DNA in host renal tissue and shows the abundance (∼50%) of *P. knowlesi* infections along with *P. vivax* (∼88%) in malaria associated AKI from our setting.

The local as well as global impact of this study is significant. In the context of India, this is the first report of *P. knowlesi* associated complications in the country and is important to highlight for better clinical diagnosis and management. In the global context, it shows the spread of *P. knowlesi* infections into regions beyond S. E. Asia and provides impetus towards urgent research that is required for understanding of *P. knowlesi* biology. It also addresses the question of *P. vivax* virulence showing that specific strains of *P. vivax* may be capable of causing severe malaria as evidenced by AKI arising out of *P. vivax*-only infections in the present study. It is a step towards understanding pathogenesis of *P. vivax* and *P. knowlesi* since we clearly demonstrate the presence of parasite derived components in host renal tissue.

Needless to say, the current study has enormous implications for public health management. Currently, in India, and several countries *P. knowlesi* is not included in preliminary diagnoses based on the belief that it does not exist in these regions. We demonstrate that this is not the case. *P knowlesi* is in circulation within the Indian population and is capable of causing malaria associated severity. In addition, in current times, there is a tremendous amount of mobility within populations and distant geographical regions. Therefore, pathogens too have the ability to travel through their hosts and reservoirs across political borders and this has been shown to happen to several pathogens including malaria. In fact resurgence of malaria in several places is attributed to international travel by their human hosts [29]. Therefore, we recommend that *P. knowlesi* must be included in preliminary screening of patients suspected to have malaria in India, as well as in all those regions where malaria is endemic and where the reservoir of *P. knowlesi*: *Macaca mulatta* has a home. Further, this study shows the need to develop advanced diagnostic methods in order to detect these pathogens with greater sensitivity and specificity since neither microscopy nor current designs of the lateral flow based rapid diagnostic tests(RDT)have the ability to discriminate between all the human malaria species, which now includes *P. knowlesi*.

## Funding

P.A. thanks SERB [Grant no. ECR/2016/000833] for funding. A.P. thanks DBT, C.K. thanks CSIR and SERB.

## Declarations

All authors declare no conflict of interest.

## Consent for publication

All authors consent towards the publication for this manuscript.

## Competing interests

The authors declare no competinginterest

## Authors’ contributions

P.A. conceived and designed the study, analyzed data, acquired funding and wrote the manuscript. AP, CK, RM and KK performed the experiments, analyzed data, prepared tables and figures. AS, GS analyzed data, provided the biopsy samples, designed study, participated in manuscript drafting. JS, AD, AB, SKA analyzed data, provided inputs into manuscript writing and were involved in the clinical aspects of patient diagnosis and management.

## Acknowledgements

All authors thank study participants

## Availability of data and materials

The dataset supporting the conclusions of this article is included within the article and as supplementary files.

