## Supplementary material for "Renal Localization of *Plasmodium vivax* and Zoonotic Monkey Parasite *Plasmodium knowlesi* Derived Components in Malaria Associated Acute Kidney Injury": Manuscript supply data (from another version of this preprint: version 2, 2019-03-13)

**Supplementary File**

### DETAILS OF METHODS

#### Selection of AKI Cases:

Archived FFPE kidney tissue blocks were retrieved for patients presenting with AKI associated with suspected or confirmed malaria during 2011-2018. Patients had presented with abdominal pain, fever and low urine output with suspected or confirmed malaria diagnosis. Kidney biopsies from cases of AKI showing acute tubular injury or necrosis (ATN) and/or acute cortical necrosis (ACN) without clinical, laboratory, or histopathological evidence of malaria were taken as controls. Initial evaluation of patients included urinalysis, ultrasound, complete blood counts and levels of creatinine, urea, electrolytes, pH and bicarbonate. Renal biopsy was performed in patients with unexplained AKI or those requiring prolonged dialysis. Diagnosis of malaria was based on peripheral blood smears. Renal biopsies were processed for light microscopy by standard techniques. Histological sections were stained with hematoxylin, periodic acid Schiff (PAS) and Jones methenamine silver stains. The glomeruli, tubules, interstitium, and blood vessels were examined in all the cores.

**Extraction of DNA from FFPE and fresh Kidney Biopsies:** Tissue of about 4 µm was scraped from FFPE blocks and transferred in 1.5ml microcentrifuge tubes and vortexed briefly after addition of 1 ml of 100 % xylene. It was centrifuged at 12,000 rpm for 1 min at 25ºC followed by heating at 50ºC for 10 mins. This step has been repeated at least 3-4 times. After all paraffin removal, 1ml of 100%, ethanol was added for washing and was repeated twice. Then ethanol was pipetted out and the tubes were dried overnight. DNA extraction was carried out using the QiagenQiAmp DNA FFPE tissue kit as per the manufacturer's protocol.

**Detection of Plasmodium species:** PCR with species specific primers for all the five *Plasmodium* species (*P. falciparum*, *P. vivax*, *P. knowlesi*, *P. malariae* and *P. ovale*) were carried out as described in Supplementary methods. The PCR amplicons were run on 1.5% gel for identification of specific bands against 50bp ladder, *P. falciparum* 205 bp, *P. vivax* 120 bp, *P. malariae* 144 bp, *P. ovale* 800 bp, and *P. knowlesi* at 207 bp). All PCR products were sequenced to confirm the presence of the correct parasite species. The PCRs were carried out in all the samples in a blinded fashion where the identity of the Disease and Control samples was revealed only after the PCRs were carried out and the results were tabulated in (Table 1; main text).

PCR conditions for typing of FFPE tissue DNA for *Plasmodium* species: The PCR conditions were standardized at initial denaturation is 95$^0$C for 5 mins, followed by 30 cycles of denaturation at 95$^0$C for 30 secs, 58$^0$C or 57$^0$C respectively for *Plasmodium falciparum* (Pf) *Plasmodium vivax* (Pv) *Plasmodium malariae* (Pm), *Plasmodium ovale* (Po) and *Plasmodium Knowlesi* (Pk) for 30secs, 72$^0$C for 30 secs, final extension at 72$^0$C for 5 mins and finally infinite hold at 4$^0$C.

**Table S1:** Oligonucleotide sequences for Plasmodium species used for PCR amplification using the standardized conditions

| ***Plasmodium* species** | **Forward primer (5'-3')** | **Reverse primer (5'-3')** |
|---|---|---|
| Pf [1] | TTAAACTGGTTTGGGAAAACCAAATATATT | ACACAATGAACTCAATCATGACTACCCGTC |
| Pv[1] | CGCTTCTAGCTTAATCCACATAACTGATAC | ACTTCCAAGCCGAAGCAAAGAAAGTCCTTA |
| Pm[1] | ATAACATAGTTGTACGTTAAGAATAACCGC | AAAATTCCCATGCATAAAAAATTATACAAA |
| Po[1] | ATCTCTTTTGCTATTTTTTAGTATTGGAGA | GGAAAAGGACACATTAATTGTATCCTAGTG |
| Pk[2] | CAGAGATCCGTTCTCATGATTTCCATGG | CTRAACACCTCATGTCGTGGTAG |


1. Snounou G, Viriyakosol S, Xin Ping Zhu, et al. High sensitivity of detection of human malaria parasites by the use of nested polymerase chain reaction. Mol Biochem Parasitol **1993**; 61:315–320.

2. Lucchi NW, Poorak M, Oberstaller J, et al. A new single-step PCR assay for the detection of the zoonotic malaria parasite Plasmodium knowlesi. PLoS One **2012**; 7:e31848.

**Table S2:** Clinical details of control groups (Abbreviations: LN; lupus nephritis, LSCS; lower segment caesarean section, PS; peripheral smear)

| Lab ID | Age/Sex | Specimen | Clinical features and Laboratory investigations | Histology | PCR findings |
|---|---|---|---|---|---|
| AKI0066 | 21/F | graft nephrectomy | Graft dysfunction, history of recent parvovirus infection | ACN secondry to graft artery thrombosis | No evidence of *Plasmodium* species |
| AKI3235 | 20/M | nephrectomy | Case of LN, post-biopsy abdominal distension and fall of hemoglobin | ACN, LN class-IV+V | |
| AKI7006 | 30/F | biopsy | History of elective LSCS,anemia, thrombocytopenia, oliguria; PS-schistocytes; dengue serology-negative | ACN | |
| AKI0339 | 50/M | biopsy | Underwent rectopexy for rectal prolapse; POD3-creatinine (8.2 mg/dl), hematuria, proteinuria, on HD | ACN, TMA | |
| AKI3417 | 15/M | biopsy | Renal allograft recipient; oliguria and rising creatinine | Diffuse ACN | |

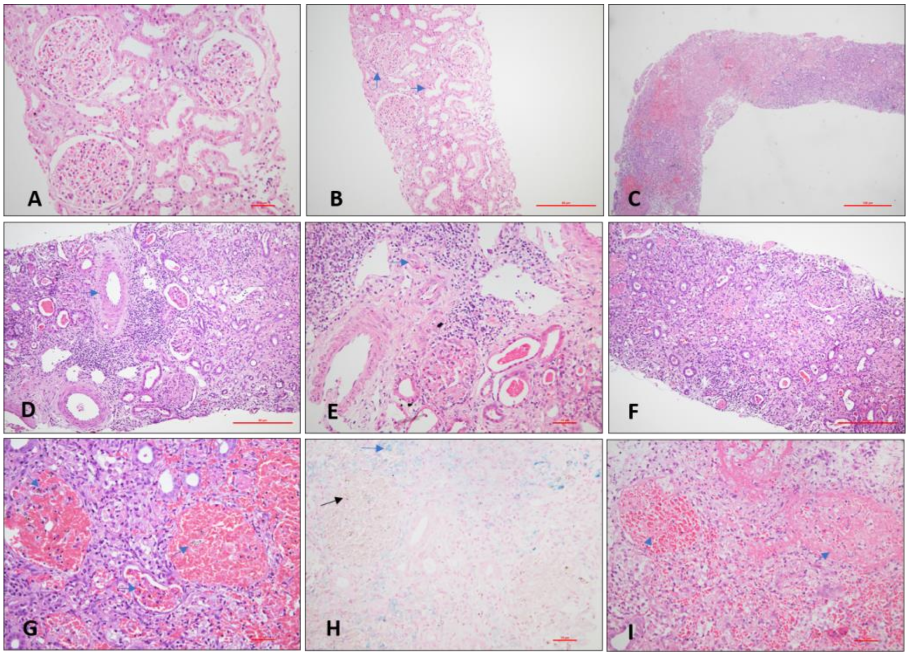


**Figure S1:** Photomicrograph depicting single linear core of renal parenchyma showing normal kidney cortex **(A)** comprising of normal glomerulus and tubules (**blue arrow**) and interstitium **(B)**. Single core of renal cortex and medulla displaying diffuse cortical necrosis containing necrosed glomeruli, tubules and blood vessels with interstitial haemorrhage involving 70% of the biopsied cortex and viable inflamed area (30%) **(C)**. Moderate mixed interstitial inflammation in the juxtamedullary area. Juxtamedullary arteries appear unremarkable **(blue arrow, D)**. Arterioles show mild medial hypertrophy with no evidence of thrombotic microangiopathy **(blue arrow, E)**. Viable cortex shows interstitial fibrosis tubular atrophy **(F)**. Glomeruli display severe mesangiolysis and congestion along with deposition of dark brown to black pigment **(blue arrow)**. Surrounding tubules shows evidence of acute tubular injury with presence of RBCs cast along with deposition of hemosiderin and blackish brown pigment in the tubular lumen and interstitium **(blue arrow, G)**. Prussian blue stain highlights presence of iron as bright blue pigment in the tubules and interstitium **(blue arrow)** while other pigment does not take the stain in the glomerulus **(black arrow, H)**. H&E slide was kept overnight in the coplin jar immersed in saturated alcoholic picric acid solution (malarial bleach), showed absence of brownish-black pigment confirming hemozoin (**black arrow**, **I)**.

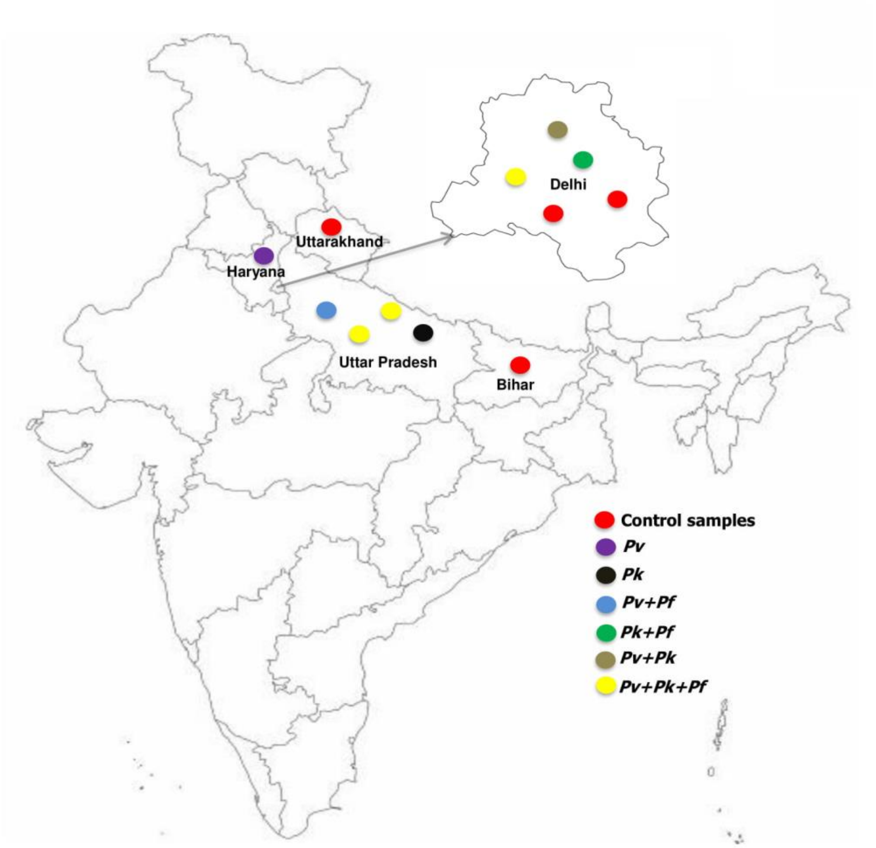


**Figure S2:** Geographic distribution of patients harboring malaria associated AKI. Location data was not available for 9 samples (3 controls, 6 cases).

### *P. knowlesi* sequencing from FFPE samples

**AKI0447_PK_F:** CCCCCAAGACCACGCGGGGGCAACGAATGTAACGTGTTAGGAAGAAAACA CCTCAGGATGTGCCAGCATCTTTTGACTTTTATTTTGGGGGGCTGCATCTACTACCACGACATGAGGTGTT CAGAGATCCGGTCTCATGATTTCGGTGTCCGTGATG

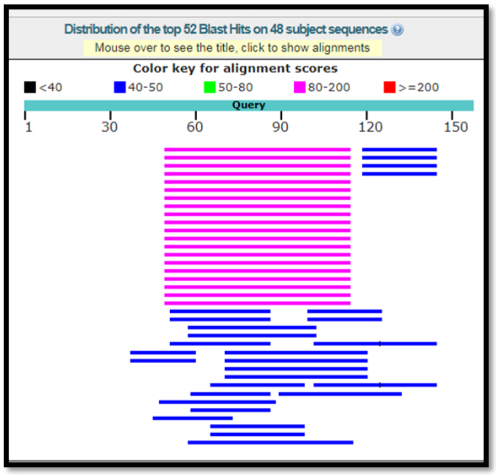


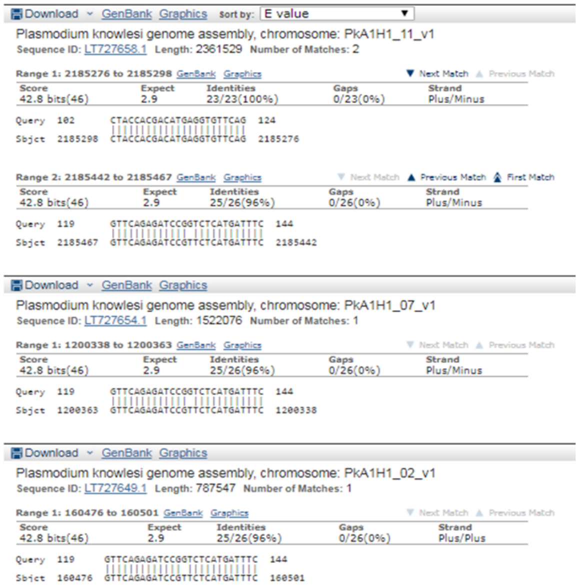


S1a: Sanger Sequencing results for *Plasmodium Knowlesi* (Pk) in kidney tissues with forward primer. Positive bands for Pk in kidney tissues of malaria infected patients, typing by PCR, was confirmed by Sanger sequencing. A nucleotide Blast (nBlast) was run with the sequencing results in NCBI (https://blast.ncbi.nlm.nih.gov/Blast.cgi) and the hits with Pk are shown in the figure.

**AKI0447_PK_R**
ACCTTAAGGAACCAGAGGAGGAGGAACAGCAGAAGAGGGAATTTCCAGCTACTAACAATTGGTGTAGTTC
ATCATCACCTGTAATGACCAAAATGTCTCCCATGGAAATCATGACAACGGATCTTTGATTTTCAAA

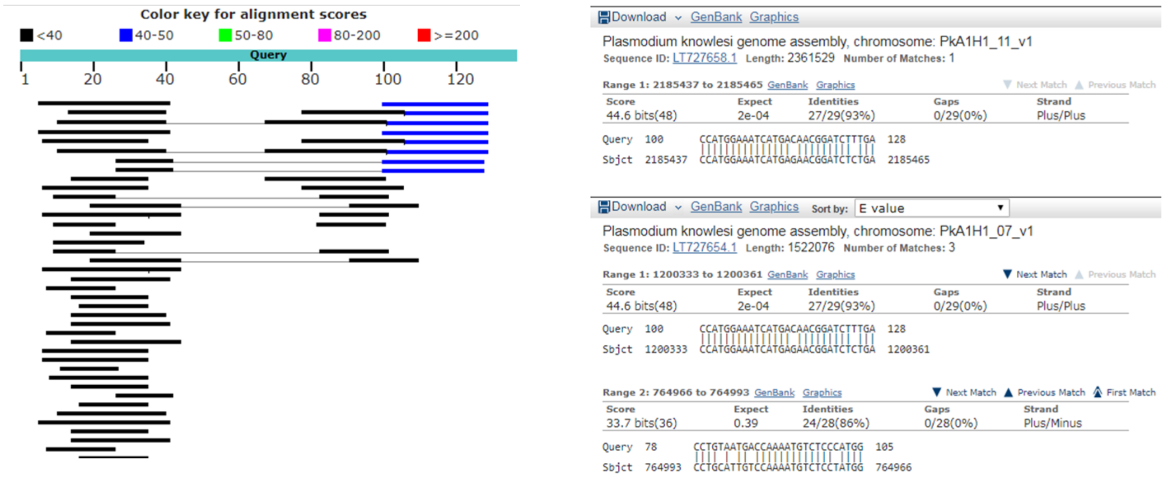


S1a: Sanger Sequencing results for *Plasmodium Knowlesi* (Pk) in kidney tissues with reverse primer. Positive bands for Pk in kidney tissues of malaria infected patients, typing by PCR, was confirmed by Sanger sequencing. A nucleotide Blast (nBlast) was run with the sequencing results in NCBI (https://blast.ncbi.nlm.nih.gov/Blast.cgi) and the hits with Pk are shown in the figure.

**AKI1056_PK_F**
CCCCCCAGGGTTGTTGTATGAAACAGAACTCTCATATTGTGAGTCGGATGAAAACAACCTCAGGAATGTGC
CAGCATCTTTTGACTTTTATTTTGGGGGGCTGCATCTACTACCACGACATGAGGTGTTCAGAGACCGG

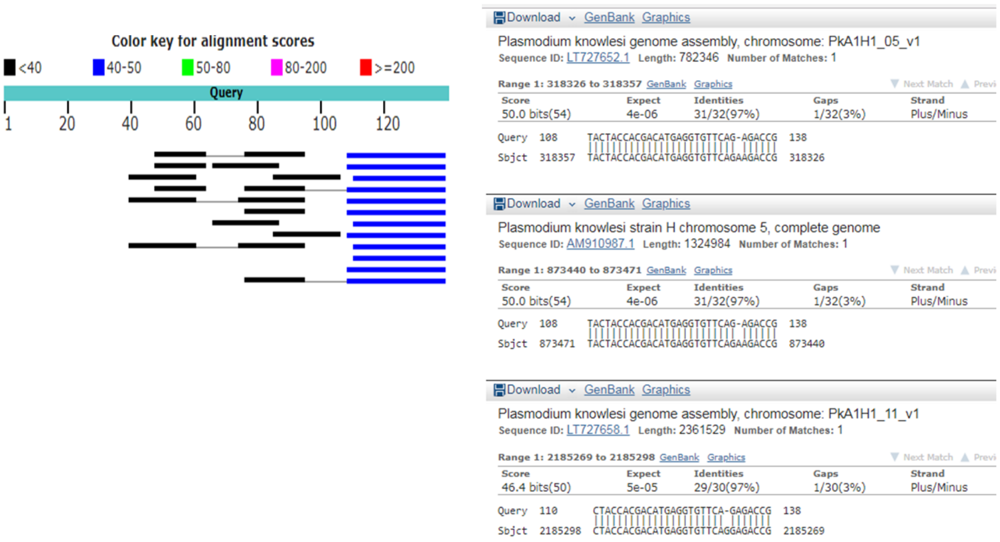


S1b: Sanger Sequencing results for *Plasmodium Knowlesi* (Pk) in kidney tissues with forward primer. Positive bands for Pk in kidney tissues of malaria infected patients, typing by PCR, was confirmed by Sanger sequencing. A nucleotide Blast (nBlast) was run with the sequencing results in NCBI (https://blast.ncbi.nlm.nih.gov/Blast.cgi) and the hits with Pk are shown in the figure.

**AKI1056_PK_R**
CGGGAGAATAGCAAAGCCACAGTTGGATGAGGGTCTTCTCAGGACTAACAATTGGTGTAGTTCATCATCAC
CTGGAATGACCAAAATGTCTCCCATGGAAATCATCACCCCGGATCTCGGATTTTCAAAT

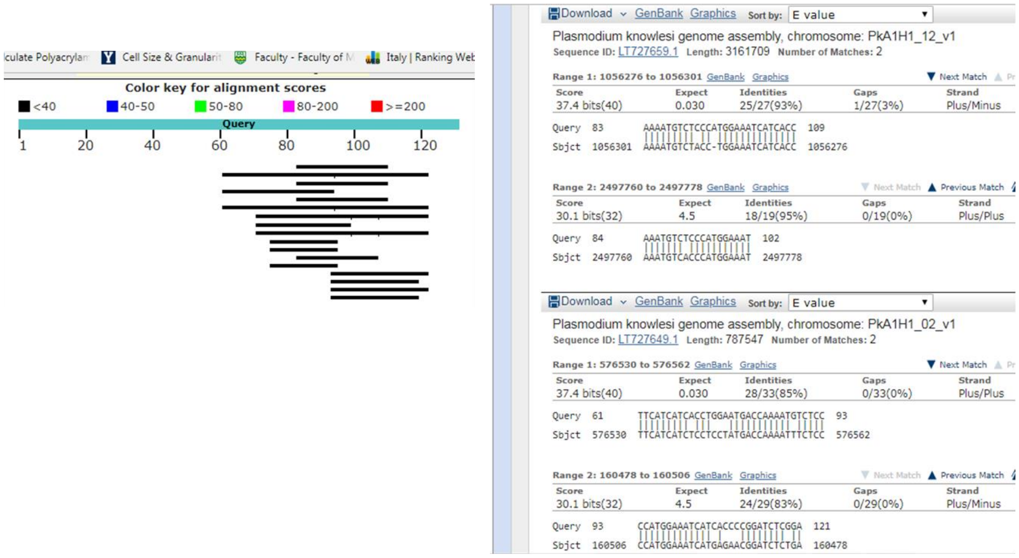


S1b: Sanger Sequencing results for *Plasmodium Knowlesi* (Pk) in kidney tissues with reverse primer. Positive bands for Pk in kidney tissues of malaria infected patients, typing by PCR, was confirmed by Sanger sequencing. A nucleotide Blast (nBlast) was run with the sequencing results in NCBI (https://blast.ncbi.nlm.nih.gov/Blast.cgi) and the hits with Pk are shown in the figure.

**AKI3873_PK_R**

CTTACCAAACCGGGGGGACCGGAGGAGGAGGAACAGAAGAAGAGGGACTTTTCAGCCACTAACAATTG
GTGTAGTTCATCATCACCTGGAATGACCAAAATGTCTCCCATGGAAATCATCACCCCGGATCTCTGATTTTC
AAA

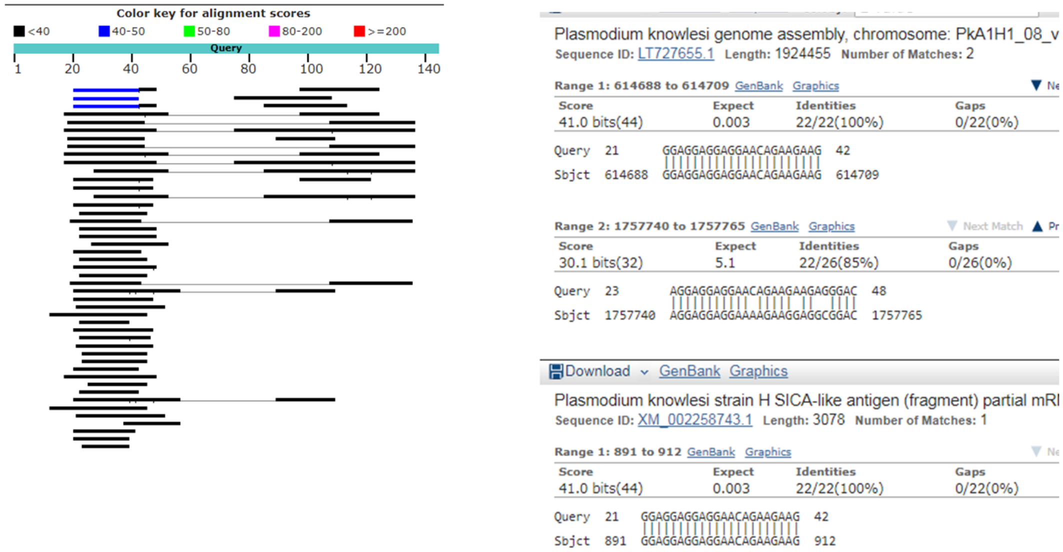


S1b: Sanger Sequencing results for *Plasmodium Knowlesi* (Pk) in kidney tissues with reverse primer. Positive bands for Pk in kidney tissues of malaria infected patients, typing by PCR, was confirmed by Sanger sequencing. A nucleotide Blast (nBlast) was run with the sequencing results in NCBI (https://blast.ncbi.nlm.nih.gov/Blast.cgi) and the hits with Pk are shown in the figure.

**AKI4424_PK_F**
TGTTGGGGTAAAGAAGGAACCTCCAGCAATTTGTTATGTGGCTGAGAAGAACCTCAGGAATGTGCCAGCA
TCTTTTGACTTTTATTTTGGGGGGCTGCATCTACTACCACGACATGAGGTGTTCAGAGATCCGTTCTCAGGA
TCCGATGTTGG

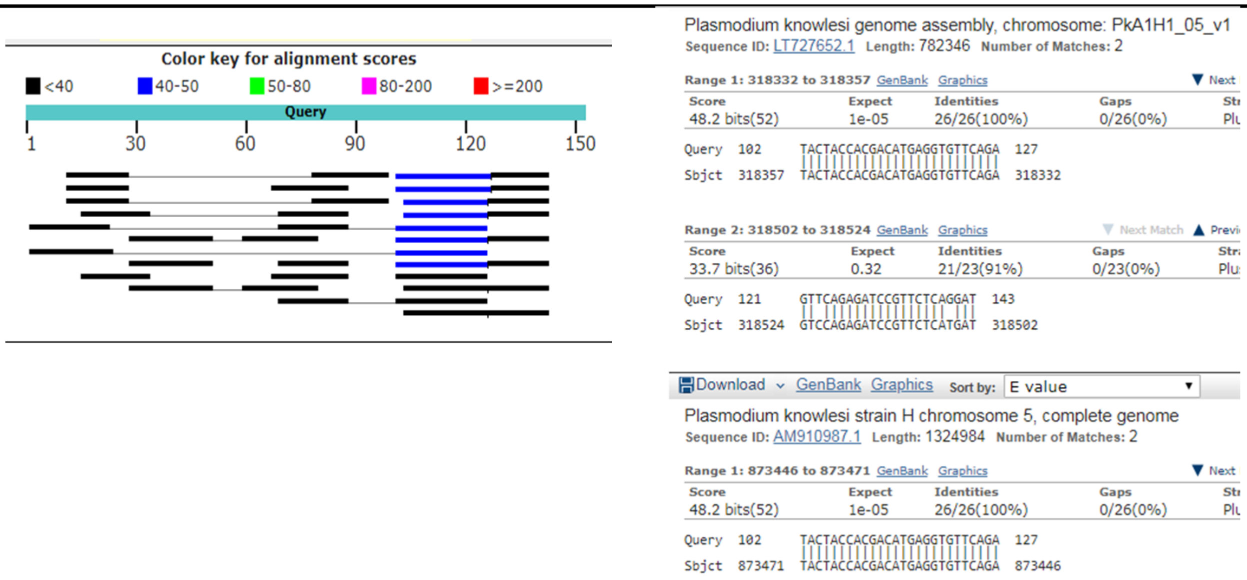


S1b: Sanger Sequencing results for *Plasmodium Knowlesi* (Pk) in kidney tissues with forward primer. Positive bands for Pk in kidney tissues of malaria infected patients, typing by PCR, was confirmed by Sanger sequencing. A nucleotide Blast (nBlast) was run with the sequencing results in NCBI (https://blast.ncbi.nlm.nih.gov/Blast.cgi) and the hits with Pk are shown in the figure.

**AKI4424_PK_R**
CTTTGAAAAACATTCCTAGAGGTTCTTTTAAGCCACTAACAATTTGGGGTAGTTCATCATCACCTGGAATGA
CCAAAATGTCTCCCATGGAAATCATGAGAACGGATCTCTGACCTCCAAAT

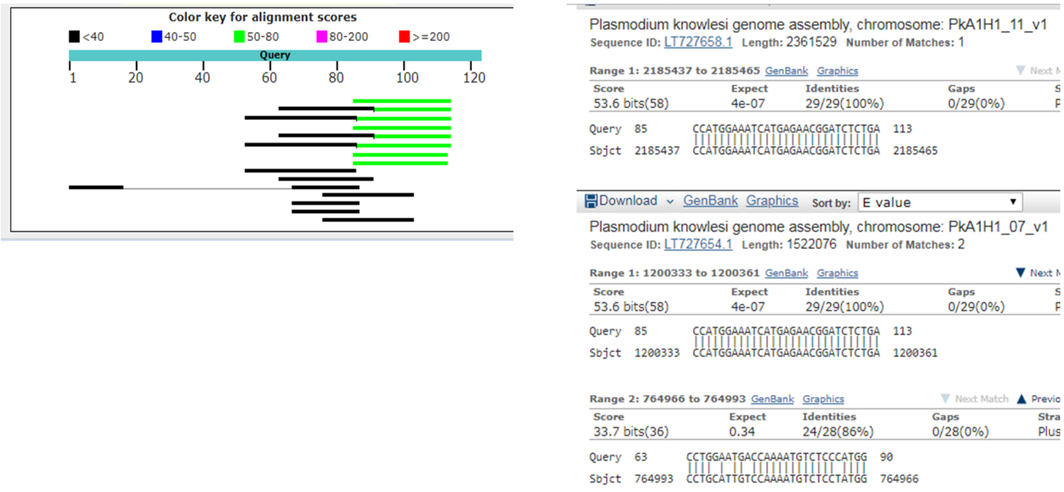


S1b: Sanger Sequencing results for *Plasmodium Knowlesi* (Pk) in kidney tissues with reverse primer. Positive bands for Pk in kidney tissues of malaria infected patients, typing by PCR, was confirmed by Sanger sequencing. A nucleotide Blast (nBlast) was run with the sequencing results in NCBI (https://blast.ncbi.nlm.nih.gov/Blast.cgi) and the hits with Pk are shown in the figure.

**AKI6691_PK_F**
CCTTCGAGGGGGGCGCGGCCGGGGGCGAAATGAGAACAGTGAGGCGGTAGAGAGAACCTCAGGAATGT
GCCAGCATCTTTTGACTTTTATTTTGGGGGGCTGCATCTACTACCACGACATGAGGTGTTTAGAGATCCGTT
CTCATGATTTCGATGGTAGTGATGCGATCCCTTCTGTGA

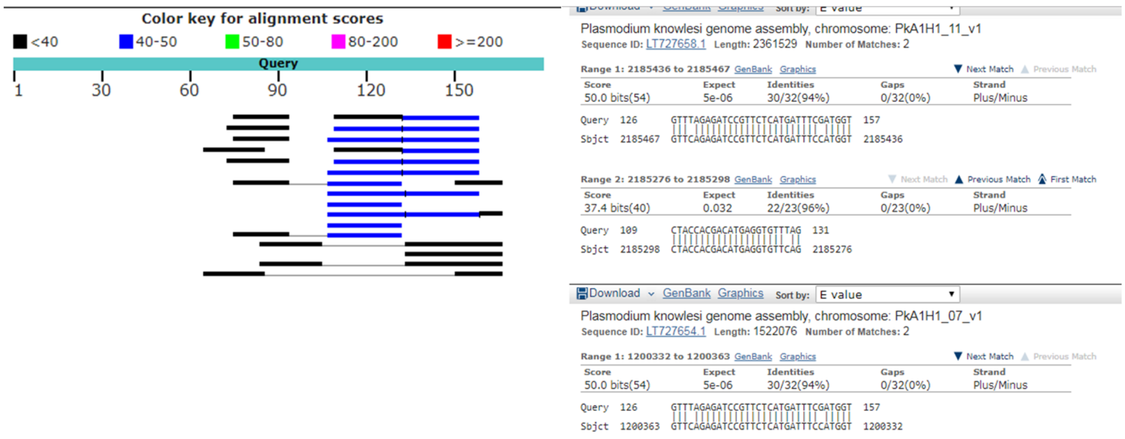


S1b: Sanger Sequencing results for *Plasmodium Knowlesi* (Pk) in kidney tissues with forward primer. Positive bands for Pk in kidney tissues of malaria infected patients, typing by PCR, was confirmed by Sanger sequencing. A nucleotide Blast (nBlast) was run with the sequencing results in NCBI (https://blast.ncbi.nlm.nih.gov/Blast.cgi) and the hits with Pk are shown in the figure.

**AKI6691_PK_R**
CTAAAGAGGGGGAGACGGCGGCGCGGGCCTGCCGATGCGGGAATTCTCAACCACAACAATCGTCGTTTTT
CTTCATCACCTGTAATGACCAAAATGTCTCCCATGGAAATCATCAGACCGGATCTCTGACCTTCAAA

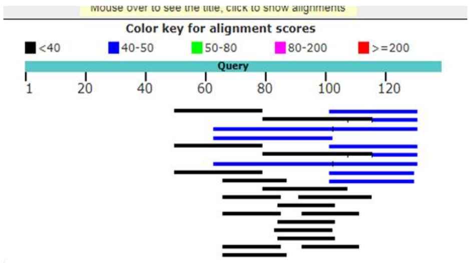


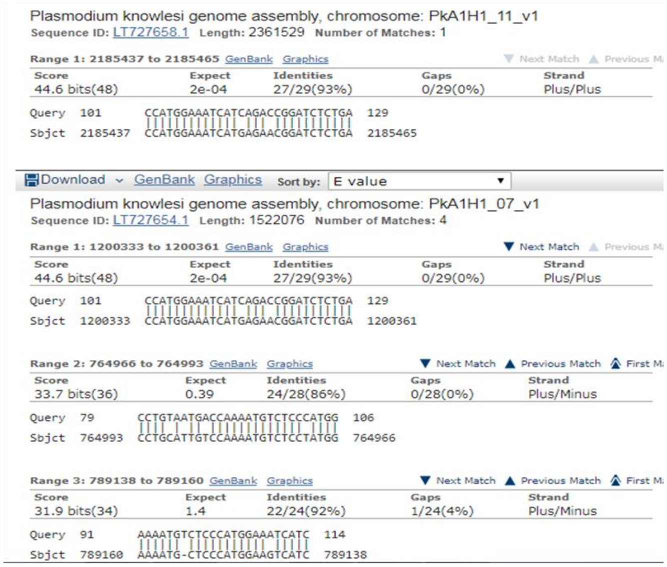


S1b: Sanger Sequencing results for *Plasmodium Knowlesi* (Pk) in kidney tissues with reverse primer. Positive bands for Pk in kidney tissues of malaria infected patients, typing by PCR, was confirmed by Sanger sequencing. A nucleotide Blast (nBlast) was run with the sequencing results in NCBI (https://blast.ncbi.nlm.nih.gov/Blast.cgi) and the hits with Pk are shown in the figure.

**AKI7167_PK_F**
TCAAAGAGCAGAGGAGAGCATAACGATGGTTAAGACTACACAGATTGTGAGTGGCTGAGAAGAACCTCAGGAATGTGCCAGCATCTTTTGACTTTTATTTTGGGGGGCTGCATCTACTACCATGGATCTCATGAGAACGGATCTCTGA

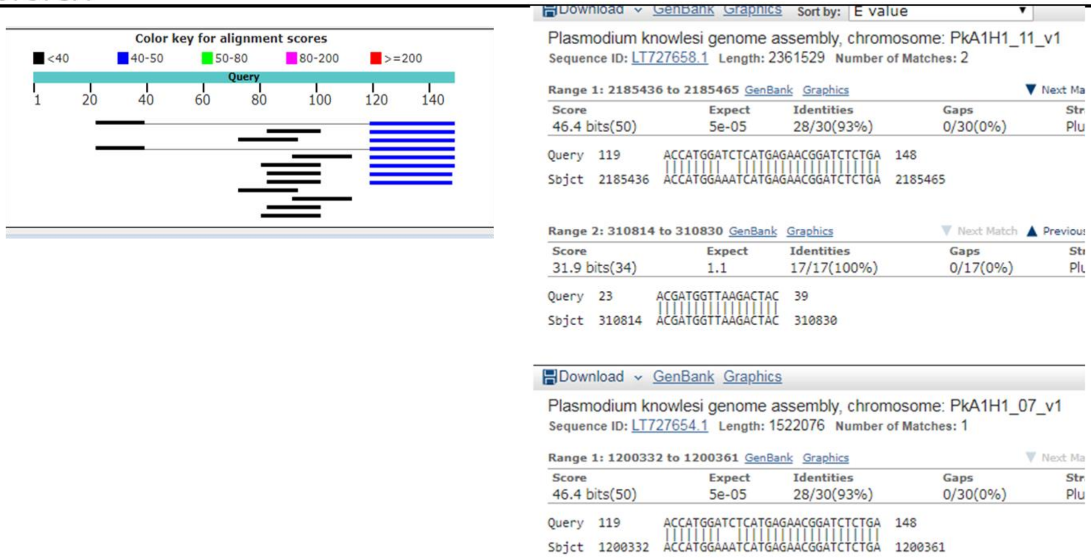


S1b: Sanger Sequencing results for *Plasmodium Knowlesi* (Pk) in kidney tissues with forward primer. Positive bands for Pk in kidney tissues of malaria infected patients, typing by PCR, was confirmed by Sanger sequencing. A nucleotide Blast (nBlast) was run with the sequencing results in NCBI (https://blast.ncbi.nlm.nih.gov/Blast.cgi) and the hits with Pk are shown in the figure.

**AKI7167_PK_R**
CTCACTATTATCGGCTCACGAGGGATTGAATCTTTGGACACTGATTCCCAGA

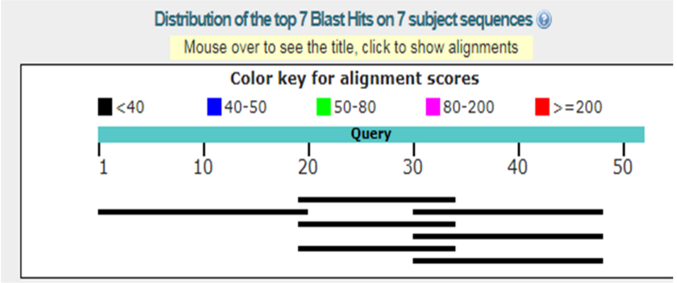


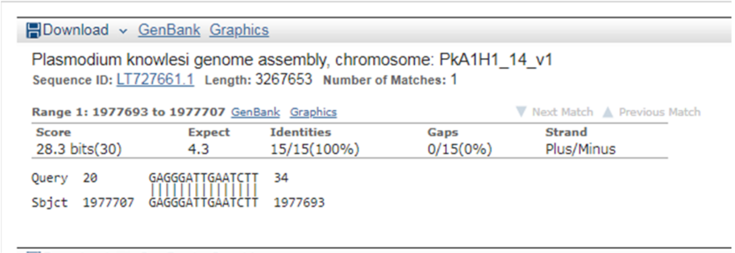


S1b: Sanger Sequencing results for *Plasmodium Knowlesi* (Pk) in kidney tissues with reverse primer. Positive bands for Pk in kidney tissues of malaria infected patients, typing by PCR, was confirmed by Sanger sequencing. A nucleotide Blast (nBlast) was run with the sequencing results in NCBI (https://blast.ncbi.nlm.nih.gov/Blast.cgi) and the hits with Pk are shown in the figure.

**AKI8299_PK_F**

CTGTCGTAATAAATAGAAGAACAGAGACAAATGAGCCGGATGAAAAGAATTCAGGAATGTGCCAGCATCTT
TTGACTTTTATTTTGGGGGGCTGCATCTACTACCACGACATGAGGTGTTCAGAGATCCGTTCTCATGATTTCC
ATGCAACTCATGAGATCTCATCTCCGATG

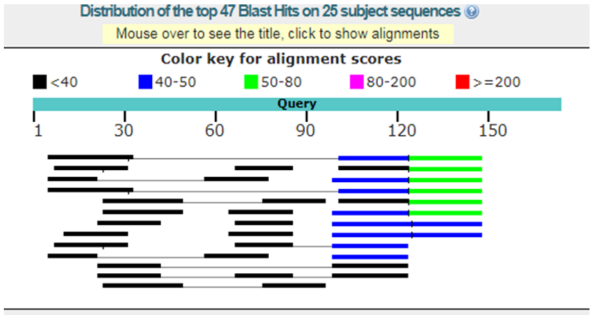


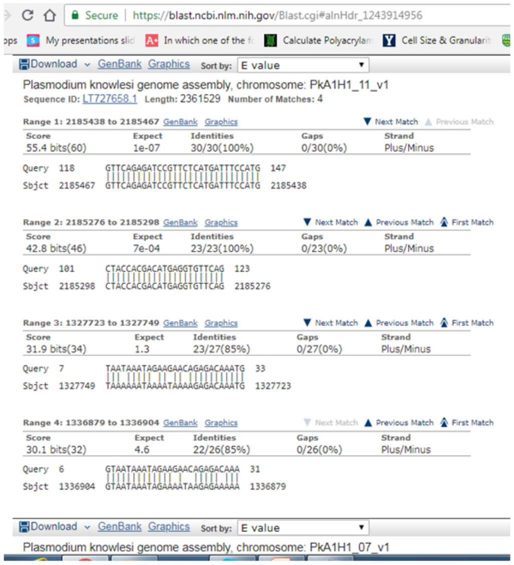


S1b: Sanger Sequencing results for *Plasmodium Knowlesi* (Pk) in kidney tissues with forward primer. Positive bands for Pk in kidney tissues of malaria infected patients, typing by PCR, was confirmed by Sanger sequencing. A nucleotide Blast (nBlast) was run with the sequencing results in NCBI (https://blast.ncbi.nlm.nih.gov/Blast.cgi) and the hits with Pk are shown in the figure.

**AKI8299_PK_R**
ATTTAGGGAAACATTCGTAGAGGGTTTTCTCAGCCACTAACAATTGGTGTAGTTCATCATCACCTGGAATGA
CCAAAATGTCTCCCATGGAAATCATGAGAACGGATCTCTGATTTTCAAAT

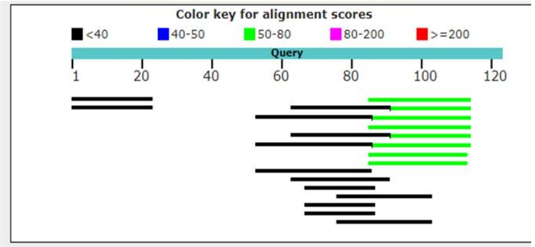


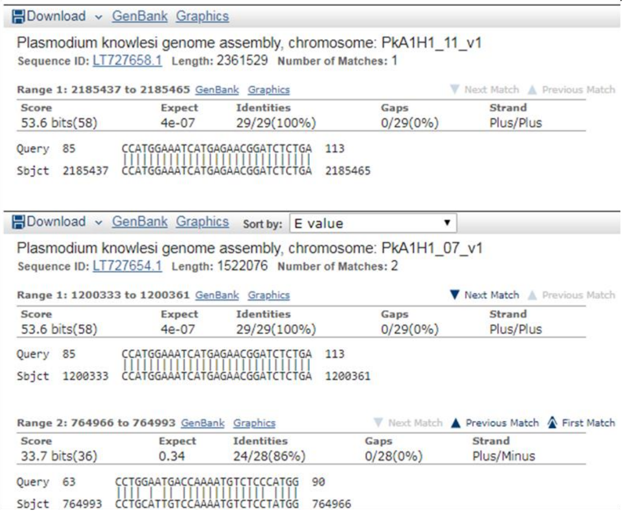


S1b: Sanger Sequencing results for *Plasmodium Knowlesi* (Pk) in kidney tissues with reverse primer. Positive bands for Pk in kidney tissues of malaria infected patients, typing by PCR, was confirmed by Sanger sequencing. A nucleotide Blast (nBlast) was run with the sequencing results in NCBI (https://blast.ncbi.nlm.nih.gov/Blast.cgi) and the hits with Pk are shown in the figure.
